# Microbial communities from lean or obese are differently shaped after apple fibres supplementation

**DOI:** 10.1101/2023.10.16.562478

**Authors:** Andrea Dell’Olio, William T. Scott, Silvia Taroncher-Ferrer, Nadia San Onofre, Jose Soriano, Josep Rubert

**Affiliations:** Laboratory of Food Quality and Design, Wageningen University & Research, Wageningen, The Netherlands; Laboratory o Human Nutrition Health, Wageningen University & Research, Wageningen, The Netherlands; Metabolomics research unit, Fondazione Edmund Mach, San michele all’Adige, Italy; Laboratory of Systems and Synthetic Biology, Wageningen University & Research, Wageningen, The Netherlands; UNLOCK, Wageningen University & Research, Wageningen, The Netherlands; Lluis Alcanyis Foundation-University of Valencia, University Clinic of Nutrition, Physical Activity and Physiotherapy, Valencia, Spain; Food and Health Laboratory, Institute of Materials Science, University of Valencia, Valencia, Spain; Department of Community Nursing, Preventive Medicine and Public Health and History of Science, University of Alicante, Alicante, Spain; Joint Research Unit on Endocrinology, Nutrition and Clinical Dietetics, Health Research, Institute La Fe, University of Valencia, Valencia, Spain

**Keywords:** Microbiota, Metagenomics, Metabolomics, Metabolic modelling

## Abstract

**Background:** Obese microbial communities differ from lean ones. Previous studies have shown how dietary fiber interventions target the gut microbiome and effectively attenuate obesity-related conditions. Nevertheless, the mechanisms by which dietary fibres shape the gut microbiota are not elucidated yet. This *in-vitro* study investigated the differences between lean and obese microbiota and how they responded to dietary interventions using a multi-omics approach.

**Results:** By employing *in vitro* digestion followed by microcolonic fermentations, we exposed obese and lean microbial communities to apple as a representative complex food matrix, apple pectin as a soluble fiber, and cellulose as an insoluble fibe. Metagenomics and metabolomics data indicated that obese and lean individuals had distinct starting microbial communities and functions. After 24 hours of exposure to different feeding conditions, the diet-responsive bacteria modulated the composition and functionality of lean and obese microbial communities. In the obese, the results suggested different mechanisms among gut commensals with an opportunistic lifestyle, allowing them to maximize their energy production from substrates breakdown and produce a specific profile of gut microbial metabolites (GMMs).

**Conclusion:** At the taxonomical and functional level, our results underscore that dietary fibres shape bacterial communities differently depending on their initial microbial composition. This modulation affects the production of GMMs. Eating foods high in fiber is recognised to promotes a healthy gut microbiome. However, the same intervention can result in varying metabolic profiles depending on the microbial communities, which may affect the host differently.

## Introduction

The human gastrointestinal microbiota is a multifaceted community of microorganisms that assumes a pivotal role in human well-being and pathological conditions[1, 2]. Various factors, such as diet, lifestyle, and host genetics, exert an influence on both the composition and function of the gut microbiome [3, 4]. The composition and function of the gut microbiome are significantly influenced by dietary factors [5, 4, 6]. Prior research has demonstrated that dietary interventions, possess the ability to influence the composition and functionality of the gut microbiome, thereby enhancing metabolic well-being [5, 7].The consumption of dietary fibers has been demonstrated to influence the gut microbiome, resulting in alterations in both the microbial composition and function that promote a more favorable phenotype[5, 6]. In particular, research has indicated that dietary interventions aimed at modulating the gut microbiome can be appealing for addressing obesity in individuals with an imbalanced gut microbiome [6, 8, 7]. At the host level, the existing body of research has established a connection between obesity and metabolic alterations that are linked to an identifiable metabolic signature involving branched-chain amino acids (BCAAs). This signature indicates an elevated breakdown of BCAAs and is closely associated with insulin resistance [9]. Moreover, previous studies have shown that there are significant differences in microbial diversity, composition, and metabolic functions within the gut microbiome of individuals who are lean compared to those who are obese [10, 11]. This study examines *in-vitro* the mechanisms through which dietary fibers influence the composition and functionality of microbial communities in both lean and obese phenotypes. We studied the fate of three substrates: apple as a representative complex food matrix, apple pectin as a soluble fiber, and cellulose as an insoluble fiber. In this study, we performed INFOGEST *in-vitro* digestion of the mentioned substrates and *in-vitro* batch fermentation from lean and obese pooled samples from clinically profiled individuals[12].We used pooled faecal samples which enables the examination of various experimental conditions, such as the incorporation of different dietary fibres into the controlled system [13] and to study more easily underlying mechanisms, especially compared to microbiome studies on faecal slurries of individuals. We used a mini-fermentation system as a reproducible batch model of the colon [14]. This makes it a feasible method for sizable screening that can later be expanded to *in-vivo* human/animal studies. We integrated a multi-omics approach including shallow shotgun metagenomics and targeted metabolomics to gain insight into the change in community structure, the possible metabolic pathways involved in substrate utilization, and the generated metabolites when different fibers are provided to modulate obese and lean microbial communities.

## 1 Discussion

### 1.1 Clinical profiling of participants and metagenomics profiling of donors and pooled samples

When conducting a comparative analysis of the anthropometric and diet variables by grouping the six profiled participants into two groups based on their body mass index (BMI), differences were found between the normal-weight and the overweight groups. Regarding anthropometric variables, significant differences were observed in body fat mass, percentage of body fat, percentage of lean body mass, and proteins. Regarding diet-related variables, significant differences were found in the intake of soluble fiber, saturated fatty acids (SFA), monounsaturated fatty acids (MUFA), iron, and fatty acids. Within the groups, there were no statistically significant differences, indicating how the subjects could be considered homogeneous in terms of anthropometry and diet. We compared the starting bacterial communities of lean and obese donors to their respective pools as shown in Figure 1 a) and 1 b). Donors within the same groups did not significantly differ in beta diversity compared to the pooled samples. Individual donors closely clustered with their relative pooled samples, as shown in Figure 1 a), suggesting that pools integrated the most relevant species. Prior research has indicated that employing a collection of faecal samples for *in-vitro* analysis does not yield a bacterial community with an atypical profile and functionality compared to that typically obtained from individual donors [13].When we performed Principal coordinate analysis (PCoA) based on Bray Curtis metric between lean and obese, including the treatments, in Figure 1 c), we found the overall composition was significantly different between obese and lean (R2=17% R2=17%), with treatment and time (R= 12% p = 0.05, R=38% p = 0.001] having significant effects. In Figure 1 b) obese and lean pooled sampled showed distinct microbial signatures depicted by unique dense clusters in the PCoA plot. Furthermore, in figure 1 a) the obese pooled samples show an altered Bacteroidetes:Firmicutes ratio compared to the lean samples. This signature has been considered a relevant marker of gut dysbiosis in obese patients [15].

**Figure 1.**
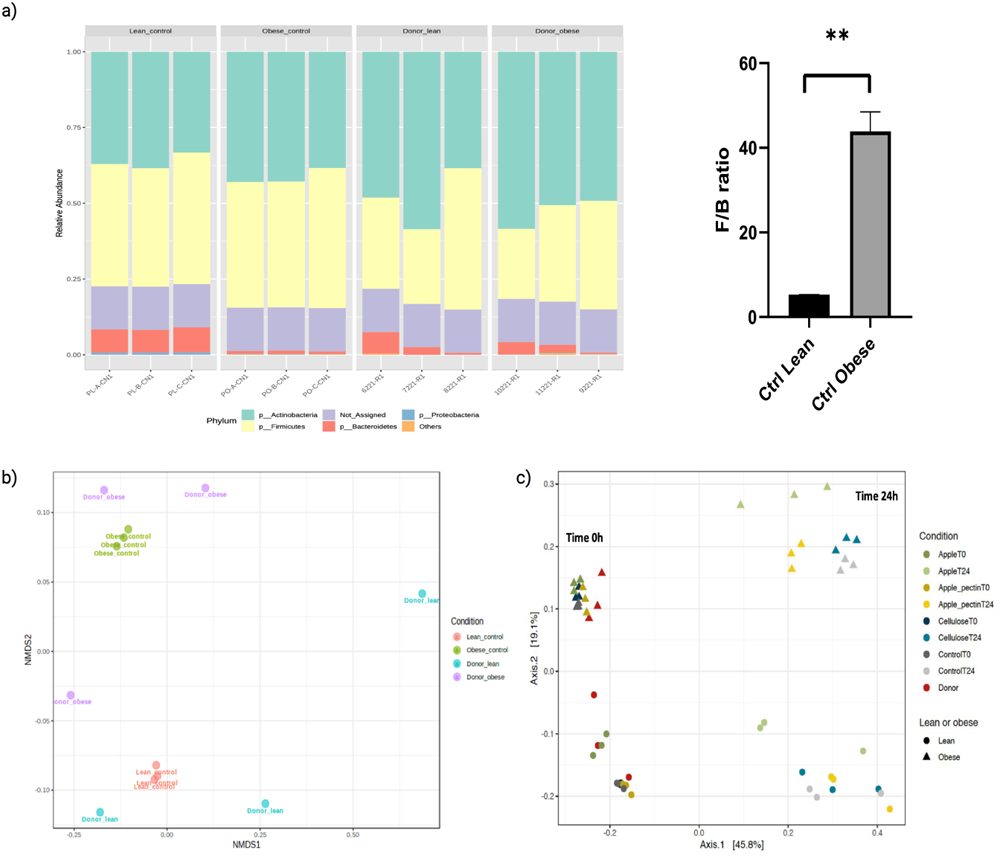
a) A graphical representation of community structure before and after pooling and the Firmicutes/Bacteroidetes ration between lean and obese donors. b) A graphical representation of community structure before and after pooling and the effect of the fermentation on the community composition.

### 1.2 Lean and obese share a contrasting metabolic signature for BCAA catabolism and hexanoic acid synthesis

When comparing the metabolic profile before and after the *in-vitro* batch culture by targeted metabolomics, there were differences between obese and lean independent of the studied treatments. The two groups differed mainly in their ability to process branched-chain amino acids and produce branched-chain fatty acids, as shown in Figure 2. In particular, the lean microbial community did not catabolize branched-chain amino acids, still resulting in the production of branched-chain fatty acids after 24 hours. By contrast, the obese microbiota resulted in the complete consumption of branched-chain amino acids and higher production of branched-chain fatty acids compared to the lean phenotype. This contrasting behaviour was independent of the given substrates. The metabolic differences in BCFAs synthesis between lean and obese phenotypes have been highlighted in recent studies and have led to contrasting opinions[16, 17, 18]

**Figure 2.**
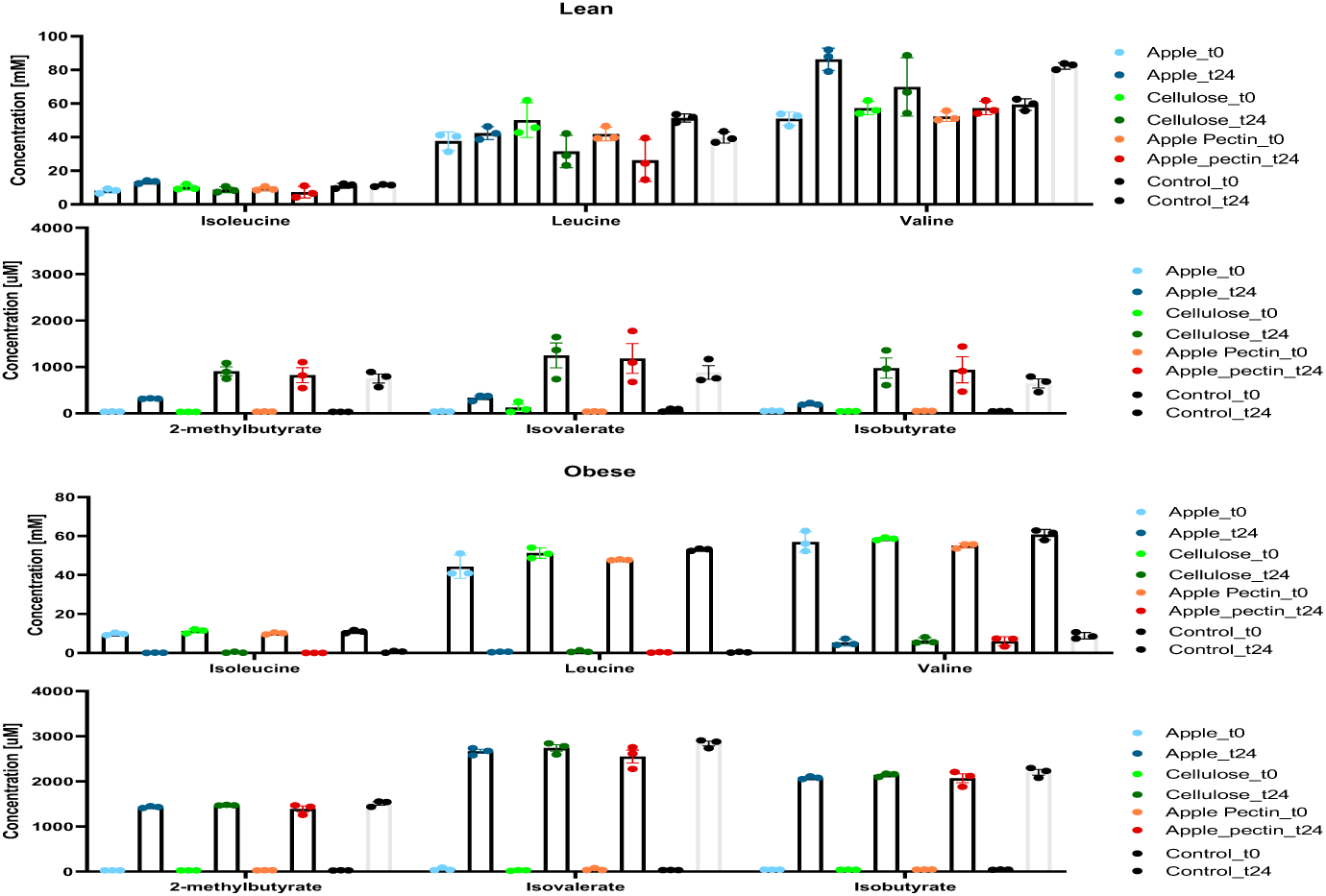
a) A graphical representation of Branched Chain Amino Acids and Branched Chain fatty acids in lean and obese before and after the treatment.

This metabolic signature seems to be a common signature of metabolic dysbiosis and could be related to diets based on high protein sources [16]. In accordance with our metabolomics data, research conducted on faecal human samples has demonstrated how the production of BCFAs is common across disease conditions. For example, obese patients exhibit elevated levels of isovaleric acid compared to individuals with a lean body mass [19]. Furthermore, individuals diagnosed with hypercholesterolemia display increased levels of isobutyric acid and isovaleric acid [19]. Additionally, patients with non-alcoholic fatty liver disease (NAFLD) have been found to have higher concentrations of isobutyric acid when compared to healthy individuals [20]. Another striking difference between the lean and obese phenotype that was treatment-independent was the synthesis of hexanoic acid. The obese microbiota consistently produced more hexanoic acid compared to the lean one as shown in 3 a). We then calculated the ratio between hexanoic acid and butyric acid after 24 hours across all the treatments and we found statistically significant differences between lean and obese in a consistent way as shown in 3 b). In our study, both BCAA catabolism and hexanoic acid synthesis signatures were independent of dietary intervention, and correlates with the previously mentioned differences in community structure between lean and obese pooled samples. These features might be considered as metabolic indicators of gut homeostais *in-vitro* between lean and obese phenotypes.

### 1.3 Diet is responsible for conferring different metabolic and taxonomic signatures in lean and obese microbiota

After supplementation with dietary fibers, significant differences were found for beta-diversity and metabolites that were strictly dependent on the type of dietary fibres provided and the starting microbial communities. As shown in figure 3 a) the metabolomic profile of the samples was highly dependent on the treatment. Both lean and obese groups showed how the type of non-digestible (ND) substrate particularly affected the functionality in terms of gut microbial metabolites (GMMs) synthesis and degradation. When the two microbial communities were exposed to apple and apple pectin, the metabolome responded differently to the intervention compared to cellulose and the control fermentation as shown in Figure 4 b). We found statistical differences between treatments for fatty acids, amino acids, and tryptophan-related metabolites. Interestingly, we detected differences in GMMs related to tryptophan metabolism in a treatment-dependent manner, especially after apple and apple pectin supplementation in the obese phenotype, as shown in Figure 4 a) and b). In particular, the apple treatment stimulated the production of Indole-3-lactic acid(ILA) and 3-Indolepropionic acid (IPA) and apple pectin stimulated Indole-3-acetic acid(IAA) and IPA production. These results highlight a tight link between dietary fibres and amino acids and their central role in shaping gut microbial metabolism and composition depending on the starting microbial community. In figure 4 b) it is evident how apple supplementation seems to have the most marked effect in both lean and obese phenotypes. Instead, apple pectin exerted a stronger effect on the obese phenotype when compared with cellulose and control samples. By looking at the species enriched by specific treatments, as shown in Figure 5 c) for obese and in Figure 9 c) for lean, apple pectin showed a bloom in *Megasphaera sp. MJR8396C*, a *Clostridiales*, after 24 hours in the obese group. The same feeding with apple pectin exerted an enrichment in *Bacteroides* species in the lean group, such as *Bacteroides ovatus* and *Bacteroides uniformis*. Apple treatment instead seemed to have a more consistent effect in both phenotypes with the enrichment of bifidobacteria together with an opportunistic enterobacteria, *Klebsiella pneumoniae*. Overall the results are summarized in Figure 5 a) and 9 a) where both metabolomics and metagenomics datasets are summarized. The data highlight how dietary intervention can cause a change in community structure and functionality depending on the type of dietary substrate and the starting microbial community. In a similar in-vitro study, Chung et al. demonstrated how the provision of apple pectin or inulin to three different human gut microbial communities from an overweight cohort led to a selective increase in the abundance of certain species[21]. Both evidence suggests that short-term dietary treatment by providing non-digestible (ND) substrates significantly impacts the overall gut microbiota *in-vitro*. Furthermore, these changes in composition depend on the initial composition of the gut microbiota [21].

**Figure 3.**
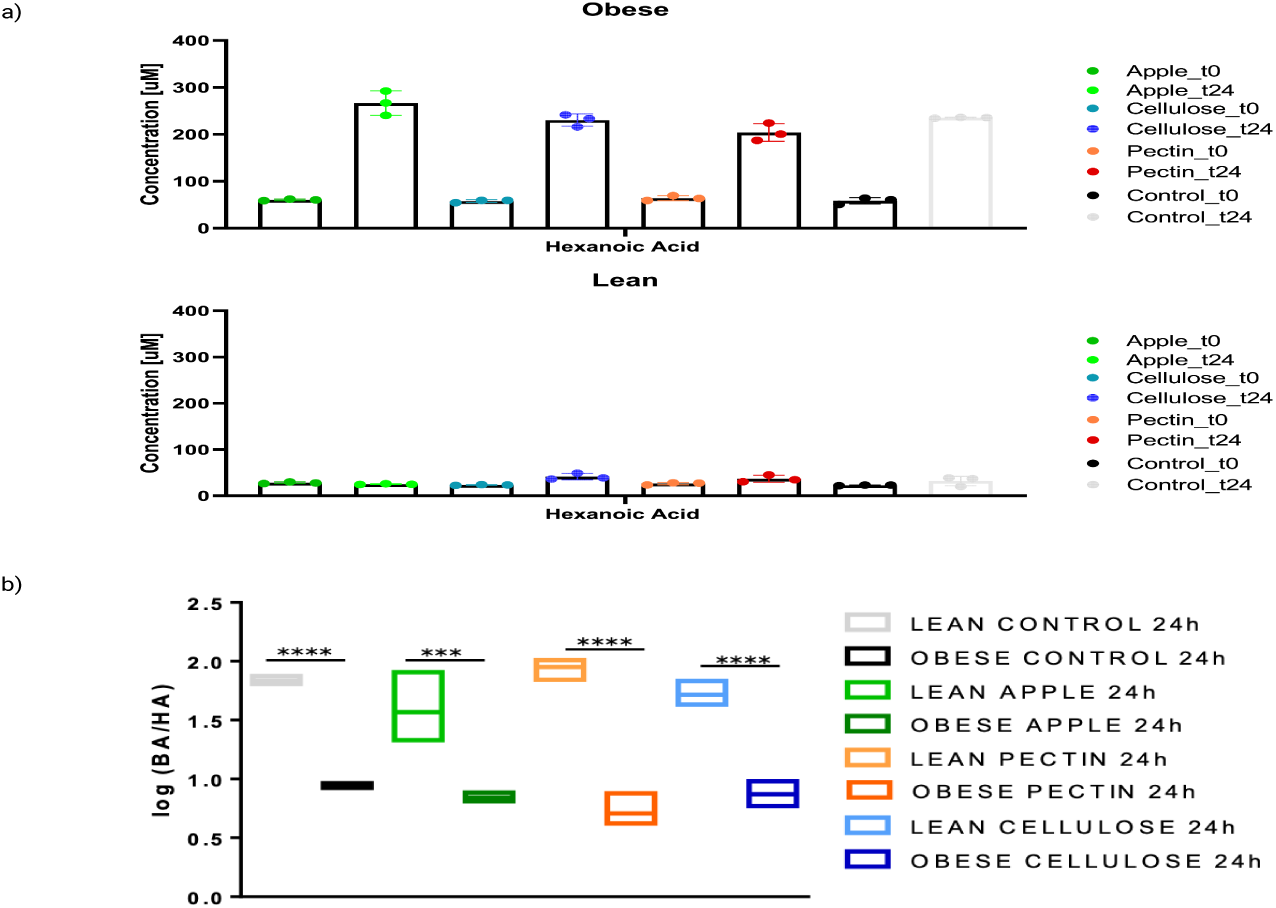
a) A graphical representation of hexanoic acid concentration across treatments b) A graphical representation of the comparison between hexanoic acid/butyric acid ratio in obese and and lean after the dieatry treatments.

**Figure 4.**
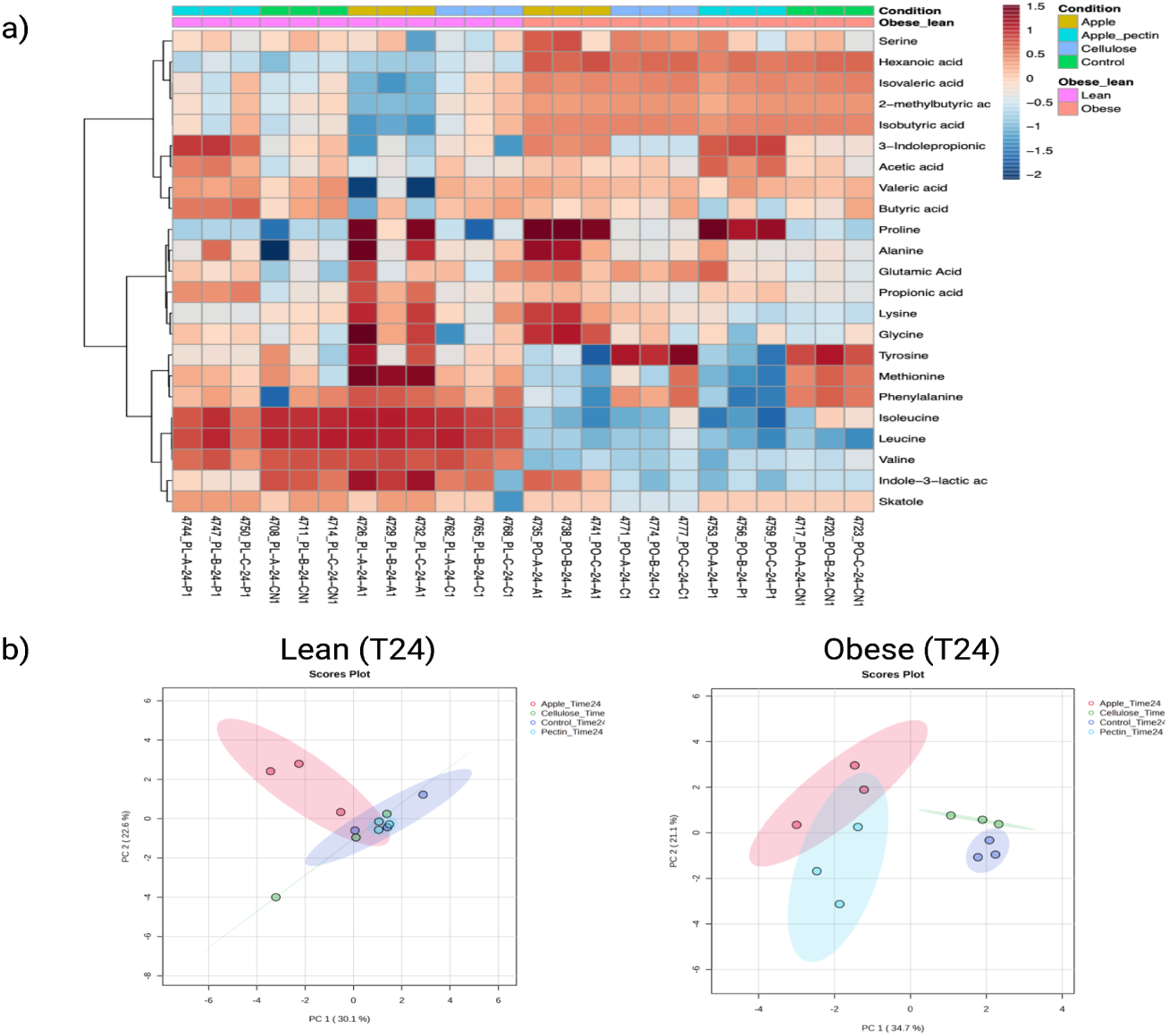
a) Heatmap of significantly different metabolites at 24 hours analyzed by two-way ANOVA corrected for condition (obese vs lean) and treatment (Apple, Apple Pectin, Cellulose, Control). b) Principal Component Analysis of targeted metabolomics after the intervention.

**Figure 5.**
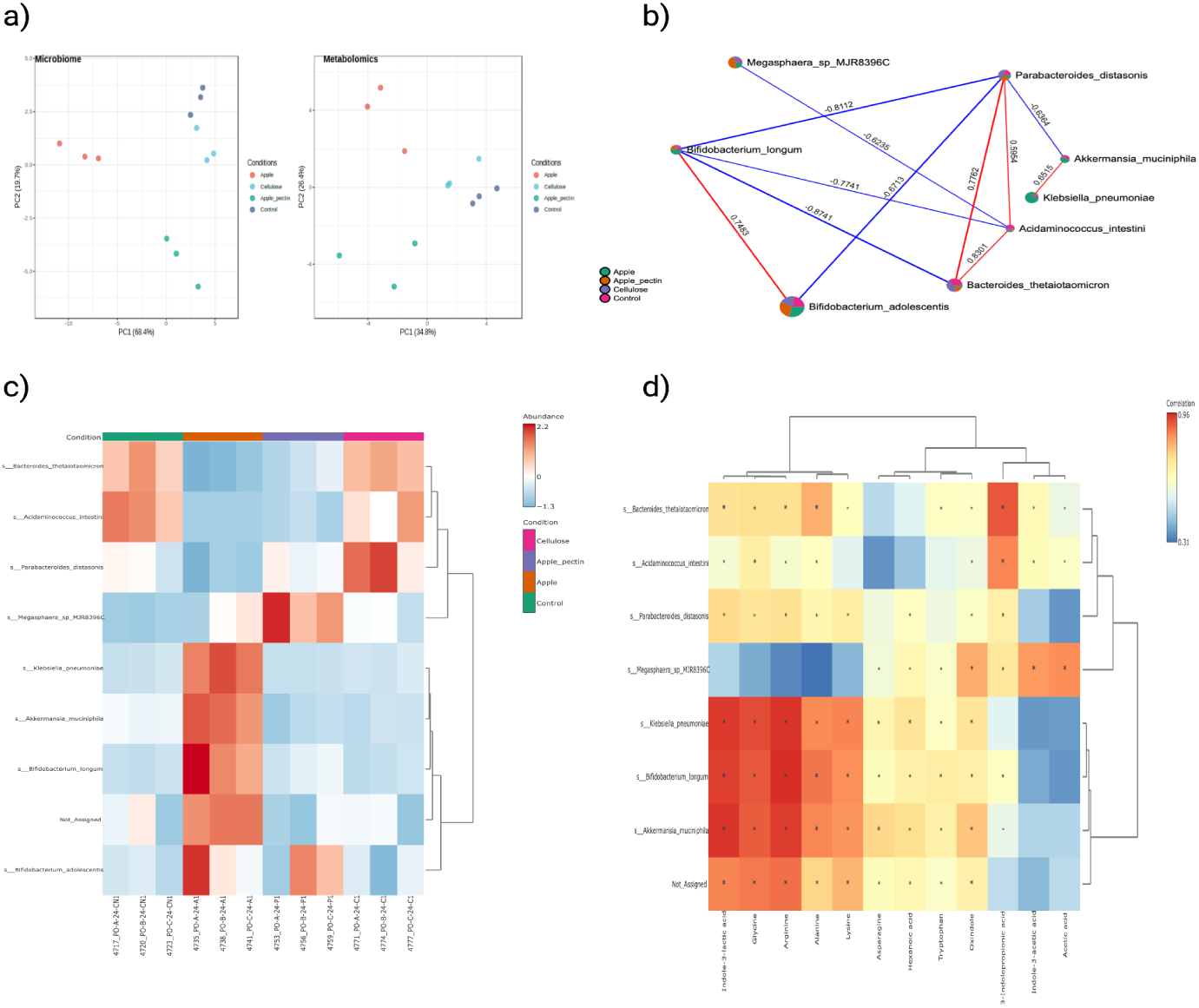
a) Comparison between metabolomics and metagenomics datasets in obese after 24 hours. b) Correlation network of significantly different species in obese at 24 hours c) Heatmap of significantly different species in obese at 24 hours analyzed by two-way ANOVA corrected for condition (obese vs lean) and treatment (Apple, Apple Pectin, Cellulose, Control) d) Correlation heatmap of significantly different species and metabolites after 24 hours of treatment in obese.

#### 1.3.1 Apple select for bacteria able to degrade phenolic acids

When looking at the scenario where apple was used as a feeding with the obese phenotype, we found a positive correlation between *Akkermansia municphila* and *Klebsiella pneumoniae* enriched together as shown in Figure 5 d) in the species correlation network. The relative abundance of these species is negatively correlated with acetic acid concentration and positively correlated with ILA concentration. To further understand the mechanisms by which apple could enrich a certain species, we looked into KEGG Orthology database to find molecular functions that were significantly enriched between the studied feedings. We selected 10 significant features with an adjusted p-value *<* 0.05 when apple feeding was compared against the other treatments. Among the significant features, of particular interest we found hydroxybenzoate decarboxylase [EC:4.1.1.61]. This enzyme belongs to the family of lyases, specifically the carboxy-lyases, which cleave carbon-carbon bonds. Interestingly, it is responsible for the non-oxidative decarboxylation of phenolic acids, suchs as p-hydroxybenzoic acid, protocatechuic acid and gallic acid [22]. These phenolic acids are one of the groups of polyphenolic compounds found in apples [23]. Phenolics in a real food matrix, such as apple, are often bound with cell wall substances such as cellulose, hemicellulose, arabinoxylans, structural proteins and pectin via ester, ether and C-C bonds [24]. This enzyme is known to be present in *Klebsiella pneumoniae* [22] genome but interestingly it is widespread in opportunistic pathogens [25]. We think therefore that the bloom of a specific opportunistic bacteria such a *Klebsiella pneumoniae* observed in both lean and obese phenotype after apple supplementation might be connected to its competitive advantage to assimilate phenolics by cleavage of C-C bonds under anaerobic conditions.

#### 1.3.2 Apple pectin select for highly specialized proteobacteria with arabinose requirements

Since we did not find significant molecular features for the apple pectin treatment and to further understand the mechanisms for the enrichment of *Megasphaera sp. MJR8396C* When apple pectin was used as a feeding, we reconstructed its metabolic genome-scale model (GSM) using CarveMe and we computed the minimal requirements for growth. Interestingly, we found a very intricate scheme with strong requirements for nitrite and nitric oxide, a simple sugar present in apple pectin, but also multiple amino acids with a preference for valine and isoleucine, and dicarboxylic acids such as glutamate and malate. The results confirm the idea of a niche specialization of Megasphaera spp. [26]. This bacteria is likely enriched when a specific combination of substrates such as simple sugars (e.g. arabinsoe), organic acids and the presence of specific amino acids are present in the gut environment *in-vitro*. We also analyzed the primary metabolism of *Megasphaera sp. MJR8396C* using GutSMASH [27]. In Figure 7, we noted the presence of a nitrogenase-like complex (Rnf) with 83% homology to the one present in *Clostridium sporogenes* and leucine reductive branch from *Clostridium diffcile*. These gene clusters indicating possible similarities with Clostridium like metabolism. In a recent study, Liu et al. discovered that *C. sporogenes* uses Rnf-based mechanism to profit from Stickland reactions [28]. *C.sporogenes* uses amino acids in Stickland pairs, coupling the oxidation of one with the reduction of another via the Stickland reaction. Notably, Liu et.al tested multiple AA pairs in *C.sporogenes* which exert different effects on the released by-products [28]. Known by-products of Stickland metabolism are, for example, IPA and IAA [28]. In the same study by Liu et al.under acetate-supplemented conditions, several substrate combinations enabled large increases in growth of *C. sporogenes* [28]. Interestingly, after 24 hours apple pectin treatment, acetic acid concentration positively correlated with *Megasphaera sp. MJR8396C*. We therefore hypothesize that acetate could be a driver for enchanced growth of an highly speialized microbe that possess traits to release GMMs such as IPA and IAA. Interestingly these molecules have been detected in the circulation of gnotobiotic mice and may also be observed in the plasma of healthy individuals [28, 29]. An increasing amount of data is therefore underscoring the importance of these metabolites in maintaining physiological balance and in the development of pathological states and according to our data there is a possibility to modulate them *in-vitro* through diet [30, 31, 29].

**Figure 6.**
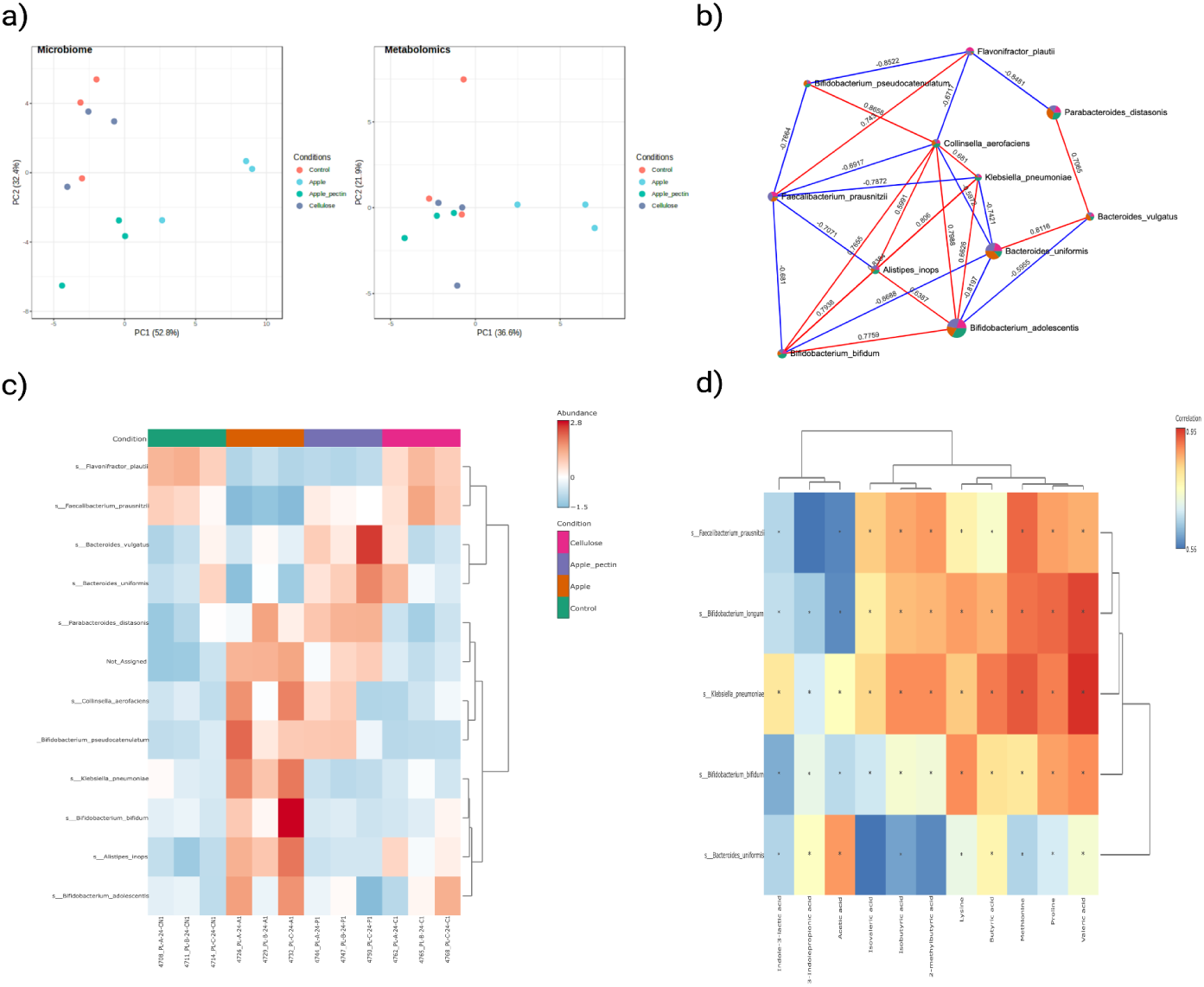
a) Comparison between metabolomics and metagenomics datasets in lean after 24 hours. b) Correlation network of significantly different species in lean at 24 hours c) Heatmap of significantly different species in lean at 24 hours analyzed by two-way ANOVA corrected for condition (obese vs lean) and treatment (Apple, Apple Pectin, Cellulose, Control).d) Correlation heatmap of significantly different species and metabolites after 24 hours of treatment in lean.

**Figure 7.**
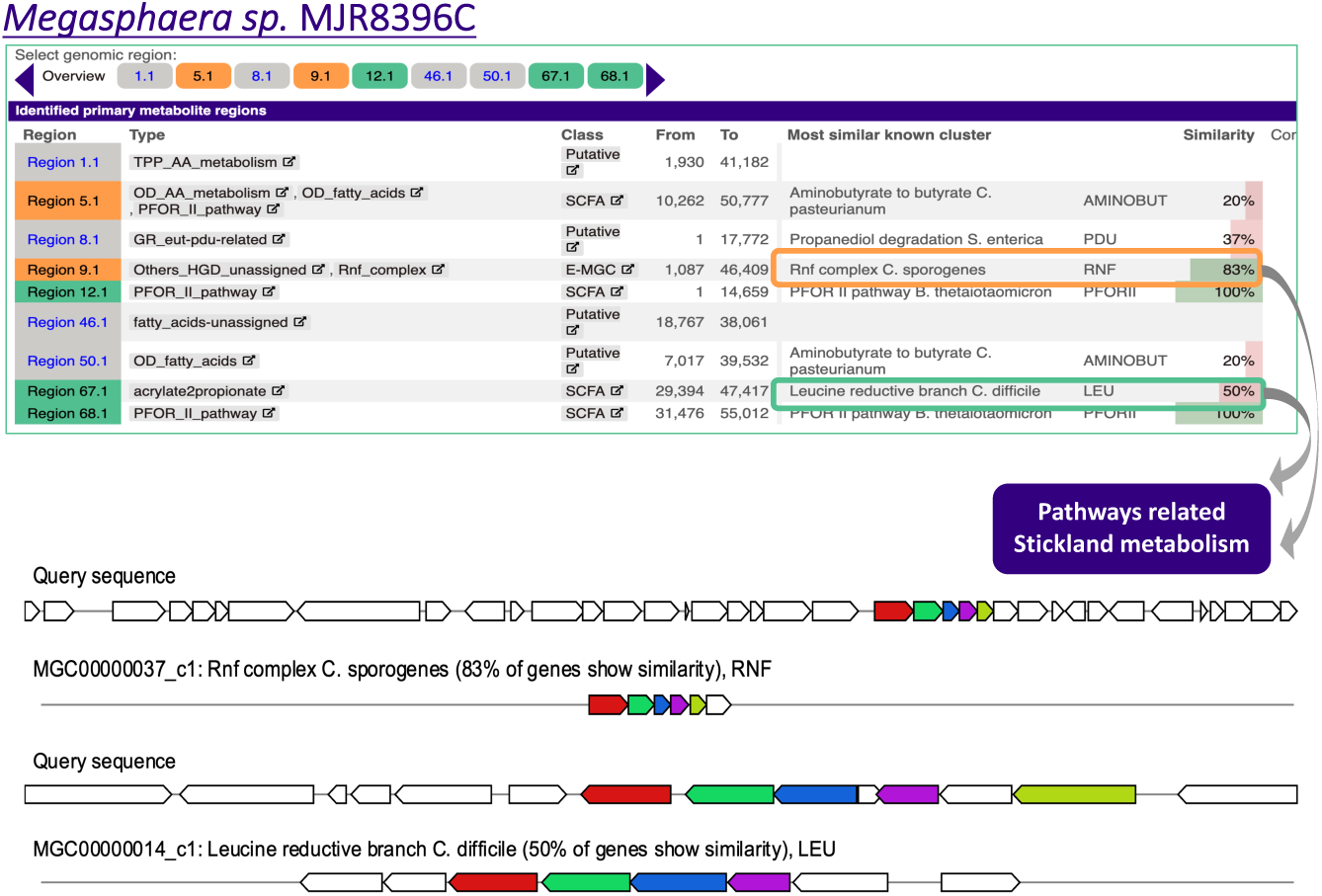
A graphical representation of GutSMASH result from the analyzed Megasphaera spMJR8396C

**Figure 8.**
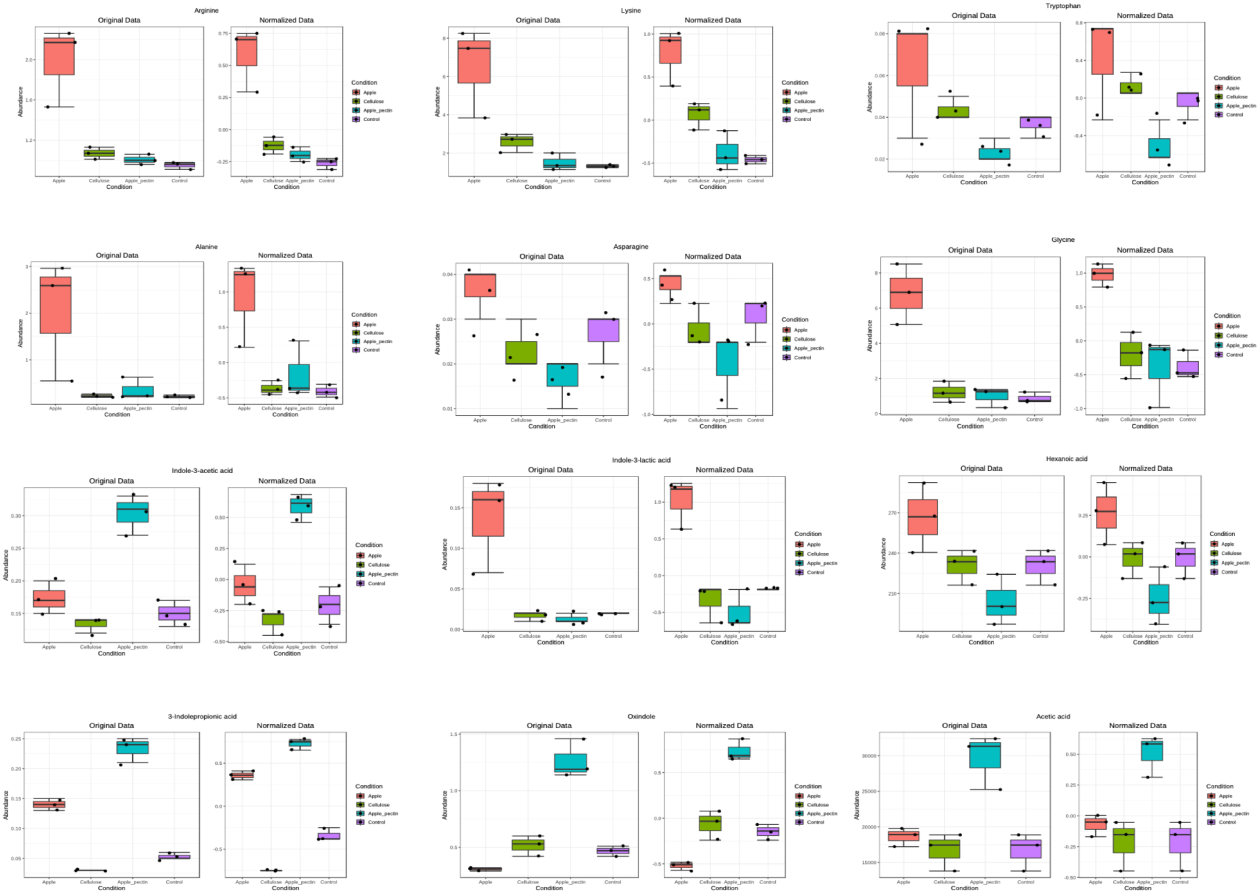
A graphical representation of metabolites that were altered in the obese phenotype depending on the dietary intervention.

**Figure 9.**
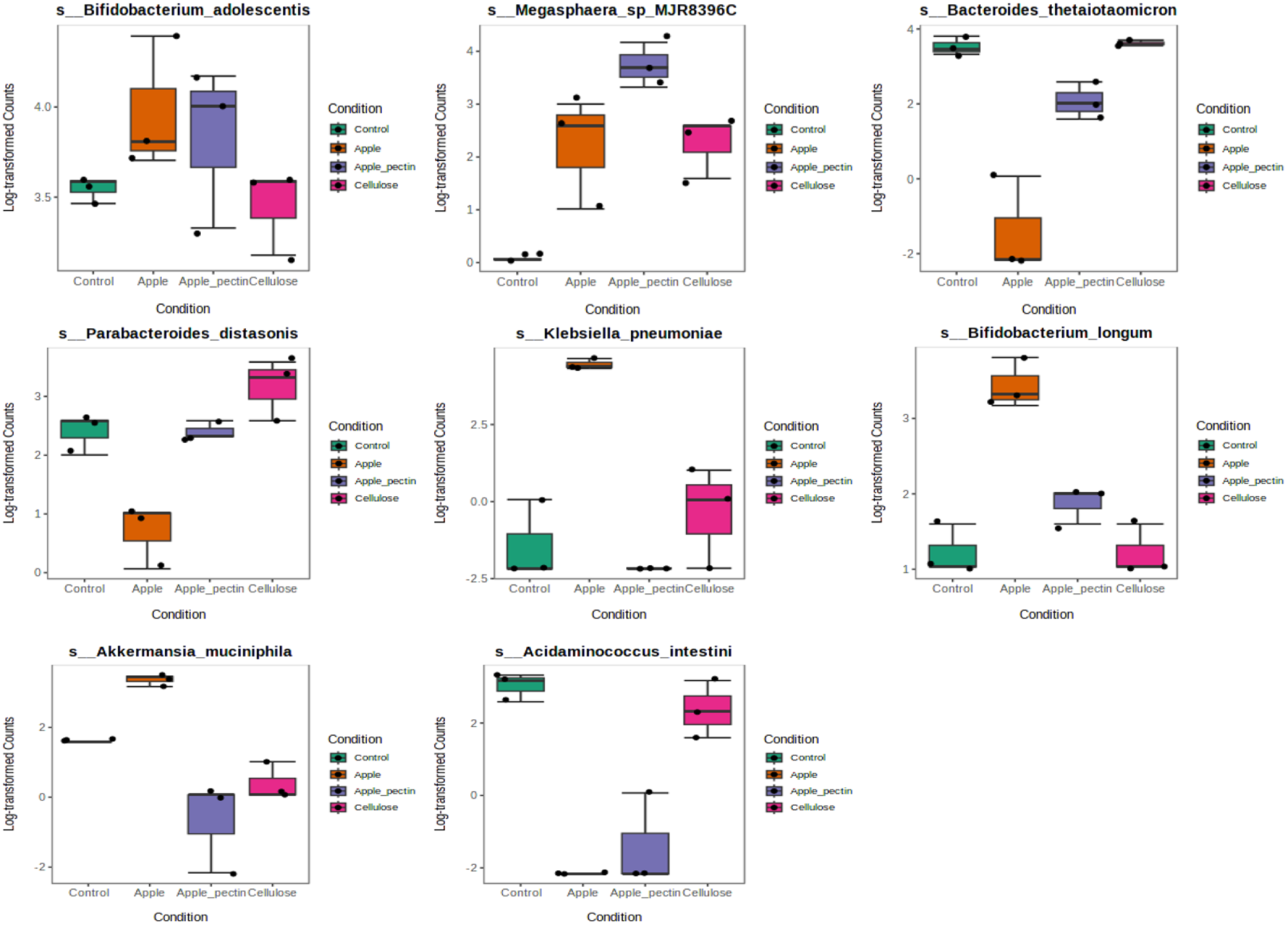
A graphical representation of species that were altered in the obese phenotype depending on the dietary intervention.

## 2 Conclusion

Dietary supplementation is gaining attention to manipulate the intestinal microbiota and alter host health. However, first, it is vital to understand how microbial strains and communities sense and degrade dietary substrates. By controlling the triangle diet-gut microbiota-host, the scientific community will open new opportunities to give nutritional recommendations that consider, for the first time, the gut-microbiome superorganisms. In this study, we employed two microbial communities, obese and lean, with significantly different microbial compositions before and after the intervention. At the functional level, the two microbial communities displayed strong metabolic differences at the starting at endpoints. The exposure of these communities to apple and apple pectin dietary substrates resulted in different functional profiles of the microbiome in terms of community structure and metabolome. We demonstrated that when the same intervention is metabolized by two different microbial communities, it can result in varying final metabolic profiles. This might potentially have a different impact on the host, locally on intestinal epithelial cells and systematically targeting different organs. By looking at the genome of the detected species, and in particular their primary metabolisms, we noticed a possible cooperation between *Akkermanisa Municiphila*, a well-known acetate producer, *Klebsiella Pneumoniae* a fastidious organism often found in disease conditions such as obesity. Furthermore, we studied in depth the metabolism of a *Clostridia* of the genre *Megasphera*, which showed complex requirements for growth and therefore a specialized enrichment during apple pectin treatment. The metabolic machinery of the detected bacteria indicated their ecological advantage to fill metabolic niches within the gut community created after exposure to the dietary fibers. This study highlights the relevance of the starting microbial community and the impact of dietary interventions at taxonomical and functional levels, describing different substrate-induced microbiota alterations that can have a substantial impact on the host. Indeed, defining gut microbial commensals as microorganisms with a positive or negative impact on the gut ecosystem is too simplistic, and context-dependent metabolisms and ecological roles depend on their microbial neighbours and substrates available. This study ultimately provides an in-depth understanding of the factors shaping nutrient hierarchies in different groups, such as obese or lean health microbial communities which can have significant implications for predicting the effects of diet on the intestinal microbiota and the resulting impact on human health.

## 3 Results

### 3.1 Clinical profiling

The study results indicate that the analyzed sample is heterogeneous regarding anthropometric and nutritional characteristics. Statistically significant differences were observed in most of the variables studied, both in anthropometric and diet-related variables.

#### 3.1.1 Comparison between normal-weight and overweight groups

When conducting a comparative analysis of the anthropometric and diet variables by grouping the participants into two groups based on their body mass index (BMI), differences were found between the normal-weight group and the overweight group:

- Regarding anthropometric variables, significant differences were observed in body fat mass, percentage of body fat, percentage of lean body mass, and proteins.
- Regarding diet-related variables, significant differences were found in soluble fiber, saturated fatty acids (SFA), monounsaturated fatty acids (MUFA), iron, C16:0, C18:0, C18:1, and C20:4.

It is important to note that the intake of C20:4 did not show significant differences in the total participants but revealed significant differences when comparing based on BMI.

#### 3.1.2 Gender comparison

The comparison of anthropometric data by gender using an ANOVA test did not show significant differences between the groups. However, when analyzing diet data by gender using an ANOVA test, significant differences were observed only for the variable B-Sitosterol, with a significance value of 0.04. No significant differences were found for the rest of the analyzed variables.

### 3.2 Beta-diversity before and after intervention

Beta diversity was assessed using principal coordinate analysis (PCoA) based on Aichison distance to quantify the compositional dissimilarity between the two groups. We performed a Permutational Multivariate Analysis of Variance (PERMANOVA) on the compositional data of treatment, group, and time point (0 or 24 hours) using Aitchison distance and Principal Coordinate Analysis (PCoA) to determine if there were significant differences based on microbiota composition. A graphical representation is provided in Figure 1. A. The results indicated that the overall composition was significantly different between obese and lean (p = 0.001), with treatment and time (P= 12%, p = 0.05, R=38%], (p = 0.001) having also significant effects. The significant difference due to treatment suggests that short-term dietary treatment had a significant impact on the overall microbiota composition.

### 3.3 Metabolites altered after dietary intervention

The provided ANOVA2 table presents significant insights into the relationship between metabolites and obesity-related conditions. Notably, Propionic acid exhibited a substantial Condition (F.val) score of 33.408 and a Condition (raw.p) value of 4.03E-07, indicating a robust association with the studied conditions. Additionally, its Interaction (F.val) score of 3.837 further suggests its potential role in influencing interactions related to obesity. Proline and 3-Indolepropionic acid also demonstrated noteworthy Condition scores of 26.129 and 22.11, respectively, emphasizing their relevance in the context of the conditions under investigation. These metabolites’ Interaction scores (4.622 and 7.181, respectively) underscore their potential impact on metabolic interactions related to obesity. In contrast, amino acids like Glycine and Tyrosine displayed higher p-values for Condition and Interaction, indicating a weaker association with the studied conditions. Furthermore, short-chain fatty acids, including Isobutyric acid and Isovaleric acid, exhibited Condition scores above 10, indicating their significant correlation with obesity-related conditions.

### 3.4 Microbial metabolites species correlations

To further understand which bacterial species might be correlated with the presence of certain GMMs in the obese phenotype we selected species of bacteria enriched by diet and correlated them with microbial signatures using MaAsLin2 (Microbiome Multivariable Association with Linear Models). Certain metabolites exhibit strong positive correlations with particular species, suggesting potential interactions within the microbiota. For instance, Glycine shows a high positive correlation of 0.908 with Akkermansia muciniphila, indicating a potential association between Glycine levels and the abundance of this species. Moreover, Indole-3-lactic acid demonstrates a strong positive correlation of 0.954 with Akkermansia muciniphila, suggesting a potential link between this metabolite and the presence of the species. In contrast, some metabolites exhibit weaker correlations with microbial species, such as Oxindole with Klebsiella pneumoniae, where the correlation coefficient is 0.723. These weaker correlations imply a less direct relationship between these metabolites and the abundance of the respective microbial species. Furthermore, metabolites like Acetic acid, Indole-3-acetic acid, and Hexanoic acid appear to have variable correlations with different species, indicating complex associations within the microbial community. Overall, this correlation table provides valuable insights into the potential connections between specific microbial metabolites and the composition of the gut microbiota, offering avenues for further investigation into the role of these metabolites in microbial ecology and host health.

### 3.5 Correlation Analysis of the detected species

In this study, we conducted a comprehensive analysis of correlations between taxonomic entities within the gut microbiota. The results reveal significant associations and interactions between these taxonomic groups, shedding light on potential ecological relationships within the microbial community. Key findings are summarized as follows: Acidaminococcus intestini and Bacteroides thetaiotaomicron: These taxonomic entities exhibited a robust positive correlation of 0.8301 (p = 8.00E-04), indicating a potential ecological interaction within the gut microbiota. This strong correlation suggests a possible co-occurrence or shared habitat niche between these taxa. Bacteroides thetaiotaomicron and Parabacteroids distasonis: A significant positive correlation of 0.7762 (p = 0.003) was observed between these two taxa, implying a potential ecological linkage between Bacteroides thetaiotaomicron and Parabacteroides distasonis. This association may be indicative of shared metabolic pathways or dependencies. Bifidobacterium adolescentis and Bifidobac-teriumlongum: These taxonomic entities displayed a strong positive correlation of 0.7483 (p = 0.0051), indicating a potential cooperative relationship or co-abundance within the gut microbiota.Akkermansia muciniphila and Klebsiella pneumoniae: A notable positive correlation of 0.6515 (p = 0.0217) was observed between these taxa. This association may suggest positive interactions or antagonistic relationships within the microbial community.Acidaminococcus intestini and Megasphaera spMJR8396C: These taxonomic entities exhibited a significant negative correlation of -0.6235 (p = 0.0303), indicating a potential competitive or mutually exclusive relationship within the gut microbiota. Akkermansia muciniphila and Parabacteroides distasonis: A noteworthy negative correlation of -0.6364 (p = 0.0261) was observed between these taxa, suggesting a potential ecological antagonism or negative interaction. Bifidobacterium adolescentis and Parabacteroides distasonis: These taxonomic entities displayed a substantial negative correlation of -0.6713 (p = 0.0168), indicative of a potential competitive or mutually exclusive relationship within the gut microbiota. Acidaminococcus intestini and Bifidobacterium longum: A strong negative correlation of -0.7741 (p = 0.0031) was observed between these taxa, highlighting a potential antagonistic or competitive interaction. Bifidobacterium longum and Parabacteroides distasonis: These taxonomic entities exhibited a significant negative correlation of -0.8112 (p = 0.0014), suggesting potential ecological interactions such as competition or exclusion. Bacteroides thetaiotaomicron and Bifidobacterium longum: These taxa displayed a striking negative correlation of -0.8741 (p = 2.00E-04), indicating a potentially antagonistic or mutually exclusive relationship.

## 4 Methods

### 4.0.1 Recruitment

The present study was conducted from November to December 2021. Six patients were profiled (four males and two females) at the University of Valencia, Clinic of Nutrition, Physical Activity, and Physiotherapy, Lluis Alcanyis Foundation. The mean age of participants was 34.8 +/-8.6 years. To each participant explained the nature and purpose of the study obtaining voluntarily informed consent from all of them. This study was approved by the Biomedical Research Ethics Committee (2018-026 – University of Trento), respecting the fundamental principles of the Declaration of Helsinki, of the Council of Europe Convention in relation to Human Rights and Biomedicine of the UNESCO Declaration. For anthropometrical assessment, participants were instructed to wear light clothing. They were positioned in accordance with the manufactures’ recommendations being height measured in a standing position and after a normal expiration using a stadiometer (SECA 225, range, 60*/*200*cm*; the precision of 1 mm, Hamburg, Germany) and weight, body mass index (BMI) (kg/m2), basal metabolic rate (kcal), fat mass (kg), body fat (%), muscle mass (kg), bone mineral mass (kg) and fat-free mass (kg) and total body water (kg) and their percentages (%)obtained using multi-frequency segmental body composition analyzer (Tanita MC780MA; Tanita Corporation, Tokyo, Japan)[32]. 24 hours before the measurements were carried out, the participants were advised to refrain from vigorous exercise, not consume any alcohol drinks, to avoid energetic drinks, and to be fasting for at least 8 h, according to [33]. For dietary consumption, the EFSA Guidance Document compiled by the EFSA Expert Group on Food Consumption Data [34], recommends that surveys cover two non-consecutive days and that the 24-hour recalls must be used for adults. The 24-hour recall interview, repeated at least once and not conducted on consecutive days, was selected as the most suitable method to get population means and distributions [35]. Estimation of portion sizes, interviewed by nutritionists/dieticians, with a picture book, including country-specific dishes, with additional household and other relevant measurements. Daily energy, macronutrient (protein, carbohydrates, and lipids), micronutrient (vitamins and minerals), and some bioactive compounds consumption was calculated using the DIAL program (Department of Nutrition UCM, Alce Ingenieria S. L., Madrid, Spain)[7].

### 4.1 Sample collection

Fecal samples were collected from obese individuals with body mass index (BMI) greater than 30 (*kg/m*) and lean (BMI lower than 25*kg/m*) individuals. Informed consent was obtained from all participants, and the institution’s ethics committee approved the study. The samples were collected using sterile containers and transported to the laboratory on ice within 2 hours of collection.

### 4.2 *In-vitro* digestion and batch fermentation

The faecal samples from obese and lean volunteers were pooled to obtain the faecal inoculum for each class. Pooling ensures that keystone microbes are not missing, which could result in compounds not being metabolized. The faecal material obtained from the three individuals per group was pooled together to obtain two fecal slurries, one from each group. The inoculum was then divided into three equal portions and exposed to apple, apple pectin, and cellulose at a concentration of 1% (w/v). The fermentation was allowed to proceed for 24 hours at 37 *◦ C*, pH = 6.9 under anaerobic conditions.

### 4.3 Metabolomics analysis

After 24 hours of fermentation, the supernatants were collected and centrifuged at 13,000 rpm for 10 minutes to remove any residual particulate matter. The supernatants were then analysed using targeted and untargeted metabolomics. Targeted metabolomics analysis on SCFAs, amino acids, and metabolites from amino acid degradation was performed using gas chromatography-mass spectrometry (GC-MS). Samples were derivatized using N-methyl-N-(trimethylsilyl) trifluoroacetamide (MSTFA) with 1% trimethylchlorosilane (TMCS) and analyzed using an Agilent 7890A GC system coupled to an Agilent 5975C MS system by the method of Lotti et al [36] . GMMs were quantified employing liquid chromatography-mass spectrometry (LC-MS) using the procedure described in. The samples were centrifuged and filtered, diluted 10-fold with Milli-Q water before being injected for targeted analysis including IPA, IAA, ILA, indole (Ind), oxindole (Oxi), indoleacrylic acid (IA), indole-3-aldehyde (I3A), tryptamine (TA), kynurenine (Kyn), and serotonin (5-HT). The metabolites were measured using a Shimadzu Nexera XR LC-20ADxr UPLC system coupled with a Shimadzu LCMS-8050 mass spectrometer (Kyoto, Japan). On a Phenomenex Kinetex 1.7 m EVO C18 100 LC column (100 2.1 mm). Chromatographic separation was carried out using 0.1 % formic acid in water (v/v) as mobile phase A and 0.1 % formic acid in methanol (v/v) as mobile phase B. By comparing the transitions and retention times with reference standards, identification was performed on a LabSolutions LCMS 5.6 (Shimadzu Corporation, Japan).

### 4.4 Metagenomics analysis

A 200 L sample of faecal slurry was subjected to automated nucleic acid extraction using the RNeasy PowerMicrobiome kit on the QIAcube device (Qiagen, Germantown, MD, United States). With the exception of skipping the phases for DNA degradation, the extraction process was carried out in accordance with the manufacturer’s instructions. The samples were bead beaten for 45 seconds using a Qiagen PowerLyzer 24 Homogenizer (Germantown, MD, United States) at a speed of 2500 rpm. In order to conduct the experiment, the samples were delivered to the QIAcube apparatus. After the extraction of the DNA, the DNA was eluted in a final volume of 100 L. Prior to sequencing, the DNA extracts were stored at a temperature of -80C. The Illumina Nextera Flex Kit for MiSeq and NextSeq platforms was used to create a pooled library. Each sample was used in this polymerase chain reaction (PCR)-based library preparation process at a rate of 1 ng. After being enzymatically fragmented and tagged with adaptors, the samples underwent PCR amplification and barcode insertion. The library was then normalised using either Illumina beads or manual techniques, and purification was completed using either columns or beads. After that, the samples were mixed and put on a plate for sequencing. On the Illumina NextSeq 550 platform, the sequencing procedure was carried out. Per sample, the experiment produced 10 million sequences. The metagenomic data were subjected to analysis using bioBakery procedures with default settings and all necessary dependencies. The raw sequence reads in this investigation were trimmed and filtered using KneadData 0.7.10. Using MetaPhlAn v 3.0, the taxonomic profiling of samples that successfully passed quality control was carried out at the species level. This programme uses alignment to a reference database of ”marker genes” to ascertain the samples’ taxonomic composition. [37]

### 4.5 Bioinformatics analysis

The metagenomics data tables were merged with the metadata, split per phenotype and time points using R. The generated datasets were analyzed using Microbiome-Analyst, an online statistical, functional and integrative analysis of microbiome data in R [38].

#### 4.5.1 Identification of significant features using multi-factory analysis

To identify significant features between the studied conditions we used MaAsLin2 (Microbiome Multivariable Association with Linear Models). This tool uses general linear models to find associations between microbial features and experimental metadata. We used an adjusted p-value cutoff of 0.05 to select significant features[39].

### 4.6 Genome mining and metabolic modelling of gut microbial species

#### 4.6.1 GutSMASH

GutSMASH was used to predict the metabolic gene clusters(MGCs) that are responsible for the synthesis of primary metabolites in gut anaerobic bacteria. The *Megasphaera sp. MJR8396C* gutsmash was generated using the assembly *GCA*001546855.1 on NCBI.

#### 4.6.2 Genome scale metabolic model reconstruction ad minimal medium calculation of Megasphaera sp. MJR8396C

The draft reconstruction and validation process involved the use of CarveMe, a tool that enables fast and automated reconstruction of genome-scale metabolic models for microbial species and communities [40]. *Megasphaera sp. MJR8396C* asssembly was available in NCBI database with NCBI RefSeq assembly GCF001546855.1. With this annotated genome sequence and the default settings of CarveMe version 1.2.2, the initial draft of *Megasphaera sp. MJR8396C* in SBML Level 3 Version 1 format was curated. Cobrapy was used to compute the minimal media requirements by providing the function minimalmedium, which obtains the medium with the lowest total import flux by default.

### 4.7 Statistical analysis

#### 4.7.1 Principal coordinate analysis (PCoA) on metagenomics data

To assess beta-diversity Principal coordinate analysis (PCoA) was performed in Microbiome Analyst 2.0 [38]. Beta diversity represents the explicit comparison of microbial communities (in-between) based on their composition. It provides a measure of the distance or dissimilarity between each sample pair. Beta diversity is calculated for every pair of samples to generate a distance or dissimilarity matrix, reflecting the dissimilarity between certain samples. The statistical methods test the strength and statistical significance of sample groupings based on the distance matrix. We used Permutational MANOVA(PERMANOVA) to test whether two or more groups of samples are significantly different based on a categorical variable found in the sample mapping file. An R value near 1 means that there is dissimilarity between the groups, while an R value near 0 indicates no significant dissimilarity between the groups. In this study Bray-Curtis dissimilarity was chosen as distance metric

### 4.8 Correlations between species based on metagenomics relative abundance

To plot correlations between species based on metagenomics relative abundance we plotted each node representing a taxon with its colour based on the user’s defined taxonomic level, and its size is based on a number of connections to that taxon. An edge connects two taxa if the correlation between the two taxa meets the p-value and correlation thresholds. The edge size also reflects the magnitude of the correlation. The blue connections represent negative correlations, while the red connections represent positive correlations. The analysis was performed in Microbiome Analyst 2.0 [38].

#### 4.8.1 Principal component analysis (PCA) on metabolomics data

Two-way ANOVA test to account for multi-factor/covariates was conducted on the metabolomics data including lean and obese samples with all the conditions studied. The test was conducted in metabonalyst using the RStatix R package, automatically determining whether Type I, II, or III errors are appropriate. The analysis was performed in Metaboanalyst Analyst[41].

#### 4.8.2 ANOVA on Anthropometric data

Descriptive analyses were conducted to calculate summary measures such as means and standard deviations for anthropometric and diet-related variables. The t-test was performed to observe statistically significant differences among the participants. In all statistical tests conducted, a significance level of 0.05 (alpha = 0.05) was set, indicating that differences with a p-value less than 0.05 were considered statistically significant. Additionally, 95% confidence intervals were calculated to estimate the precision of the mean estimates and other statistical parameters. Regarding group comparisons, an analysis of variance (ANOVA) was conducted to assess the differences in anthropometric characteristics and diet among the established groups. The groups were formed based on body mass index (BMI), differentiating between the normal-weight and overweight groups. A comparison by gender was also performed, dividing the participants into male and female groups. Furthermore, additional tests using multiple comparison tests, such as the t-test with Bonferroni correction, were conducted to evaluate significant differences in specific variables. The statistical methodology used in this study was conducted using IBM SPSS Statistics version 27 software (IBM Corp., Armonk, New York, NY, USA).

## Supporting information

Supplementary tables supporting the findings

Megasphaera_sp_MJR9396C Metabolic Model

## Acknowledgements

No acknowledgements.

## Funding

Nothing to declare.

## Availability of data and materials

Materials described in the manuscript, including all relevant raw data, will be freely available upon request.

## Ethics approval and consent to participate

The study as been approved by the ethical committee of the University of Trento with the protocol number 2018-026.

## Competing interests

The authors declare that they have no competing interests.

## Consent for publication

All the authors gave written consent before publication of the work. . . .

## Authors’ contributions

Andrea Dell’Olio performed the bioinformatics work, analyzed the data, prepared figures and/or tables, authored drafts of the paper and approved the final version. William T. Scott, J, contributed to generation of Genome Scale Models, reviewed drafts of the paper, and approved the final version. Silvia Taroncher-Ferre, performed sample collection, performed statistics on clinical data. Nadia San Onofr performed sample collection, performed statistics on clinics data. Jose Soriano performed the statistics on clinical data reviewed drafts of the paper and approved the final version. Josep Rubert conceived the work, performed the *in-vitro* fermentation and metabolomics analysis, reviewed drafts of the paper, approved the final version.

## Notes

### Competing Interest Statement

The authors have declared no competing interest.

